# Astrocytic Cathepsin D truncates α-synuclein and promotes Lewy neurite-like aggregation in neurons

**DOI:** 10.1101/2025.10.03.680233

**Authors:** Hiroshi Hanafusa, Kota Ohashi, Hiromu Shojima, Naoki Hisamoto, Takeshi Kameyama, Muneaki Miyata, Kiyohito Mizutani, Yoshimi Takai, Kunihiro Matsumoto

**Affiliations:** Division of Biological Science, Graduate School of Science, Nagoya University, Chikusa-ku, Nagoya 464-8602, Japan; Division of Pathogenetic Signaling, Department of Physiology and Cell Biology, Graduate School of Medicine, Kobe University, Kobe, Hyogo 650-0047, Japan; Division of Pathogenetic Signaling, Institute of Advanced Medical Science, Tokushima University, Tokushima, Tokushima 770-8503, Japan

## Abstract

Parkinson’s disease (PD) and related synucleinopathies are marked by the accumulation and propagation of α-synuclein (α-syn) aggregates, a process primarily studied in neurons. However, the contribution of astrocytes to this process remains unclear. Here, using a physiologically relevant neuron–astrocyte co-culture system that recapitulates tripartite synapse architecture, we show that astrocytes actively process and propagate α-syn aggregates. Astrocytes internalize α-syn pre-formed fibrils (PFFs) and cleave them into C-terminally truncated, seeding-competent species via the lysosomal protease Cathepsin D (CtsD). These truncated species are subsequently transferred to neurons, where they promote the growth of Lewy neurite (LN)-like aggregates. Notably, α-syn PFF exposure disrupts lysosomal membrane integrity in astrocytes, leading to CtsD upregulation. These findings reveal a feed-forward mechanism in which astrocytic lysosomal dysfunction amplifies α-syn pathogenicity. Our findings identify astrocytes as active contributors to α-syn propagation and highlight the astrocytic lysosomal pathway as a potential therapeutic target in PD.

## INTRODUCTION

Parkinson’s disease (PD) is the second most common neurodegenerative disorder, characterized by the progressive degeneration of dopaminergic neurons in the substantia nigra pars compacta and the accumulation of α-synuclein (α-syn)-rich inclusions, known as Lewy bodies (LBs) and Lewy neurites (LNs)^1–5^. α-syn is a presynaptic protein involved in synaptic vesicle trafficking and neurotransmitter release, though its precise physiological role remains incompletely defined^6–11^. A growing body of evidence supports the “prion-like” hypothesis, in which misfolded α-syn aggregates propagate between cells by templating the misfolding of endogenous α-syn in recipient cells^12–15^. This phenomenon can be reliably modeled by treating cultured neurons with α-syn pre-formed fibrils (PFFs), which induce time-dependent aggregation that recapitulates the structural maturation of α-syn pathology from punctate inclusions to LNs and ultimately to LB-like aggregates^16–18^.

While most studies have historically focused on neurons as both the source and target of α-syn pathology, recent findings suggest a significant role for glial cells, particularly astrocytes, in modulating α-syn proteostasis^19–23^. Astrocytes maintain neural homeostasis by providing metabolic and trophic support, clearing extracellular proteins, and shaping synaptic architecture through the formation of tripartite synapses, in which astrocytic processes intimately engage with both pre- and post-synaptic terminals^24,25^. In PD, astrocytes are capable of internalizing and degrading extracellular α-syn aggregates, a function generally considered neuroprotective^26,27^. However, chronic exposure to pathological α-syn may overwhelm this clearance capacity, triggering astrocytic dysfunction, inflammatory responses, and potential neurotoxicity^28–32^. Crucially, whether astrocytes act solely as passive scavengers or actively contribute to the processing and spread of pathogenic α-syn species remains unclear. A major limitation in addressing this issue has been the lack of physiologically relevant models that accurately recapitulate neuron–astrocyte interactions in the context of α-syn pathology. To overcome this, we recently developed a co-culture system consisting of primary hippocampal neurons and astrocytes derived from neurospheres, in which astrocytes form highly ramified processes and establish tripartite synaptic contacts with neurons, closely mimicking in vivo structural and functional relationships^33^.

Cathepsin D (CtsD), a lysosomal aspartic protease, plays a key role in α-syn degradation^34–37^. Genetic ablation of CtsD in the central nervous system of mice leads to spontaneous α-syn aggregation, suggesting its critical role in maintaining proteostasis^38^. Paradoxically, CtsD also cleaves the C-terminal region of α-syn fibrils, generating truncated forms with enhanced aggregation and seeding potential^35,39–41^. These truncated α-syn species are enriched not only in neuronal LBs but also in astrocytes in PD and dementia with Lewy bodies (DLB)^2,42–46^, suggesting a complex, context-dependent role for CtsD in disease pathology. However, it remains unknown whether astrocytic CtsD contributes to α-syn truncation and the formation of pathological aggregates in neurons.

In this study, we utilized our advanced neuron–astrocyte co-culture model to investigate whether astrocytes play an active role in processing and propagating α-syn pathology. We show that astrocytes internalize α-syn PFFs and cleave them via CtsD-dependent C-terminal truncation. The resulting truncated species are transferred to neurons, where they promote LN-like aggregation. This process is further accompanied by lysosomal membrane disruption and upregulation of CtsD in astrocytes, suggesting a feed-forward mechanism. Our findings identify astrocytic lysosomal processing as a key step in the propagation of pathogenic α-syn and highlight the astrocyte–neuron axis as a potential therapeutic target in PD.

## RESULTS

### Astrocytes promote the elongation of LN-like α-syn aggregates in neurons

To model α-syn aggregation, we applied pre-formed fibrils (PFFs) of recombinant α-syn to neuronal cultures^47^. Consistent with previous reports^17^, α-syn PFF treatment induced a time-dependent formation of Ser-129– phosphorylated α-syn (pS129–α-syn)-positive inclusions. In our system (Fig. S1a), punctate aggregates first appeared along neuronal processes by day 6 post-α-syn PFF treatment, followed by the emergence of filamentous LN-like structures by day 10 (Fig. S1b). Larger, somatic LB-like inclusions were also observed near neuronal nuclei. These aggregation patterns faithfully recapitulate key pathological features of PD^17,18^.

To investigate how astrocytes influence the progression of α-syn pathology, we utilized a physiologically relevant co-culture system comprising primary hippocampal neurons and astrocytes derived from mouse neurospheres^33^. Astrocytes were introduced into neuronal cultures at day in vitro (DIV) 10 and maintained until DIV24 (Fig. 1a). Immunolabeling with the glutamate transporter EAAT1 confirmed the identity of astrocytes, which exhibited a highly ramified morphology and extended fine processes that contacted MAP2-positive neuronal dendrites (Fig. S2a). These processes frequently localized near both pre-synaptic (VGLUT1) and post-synaptic (HOMER1) markers (Fig. S2b–d), indicating that astrocytes engaged in tripartite synapse formation.

**Fig. 1.**
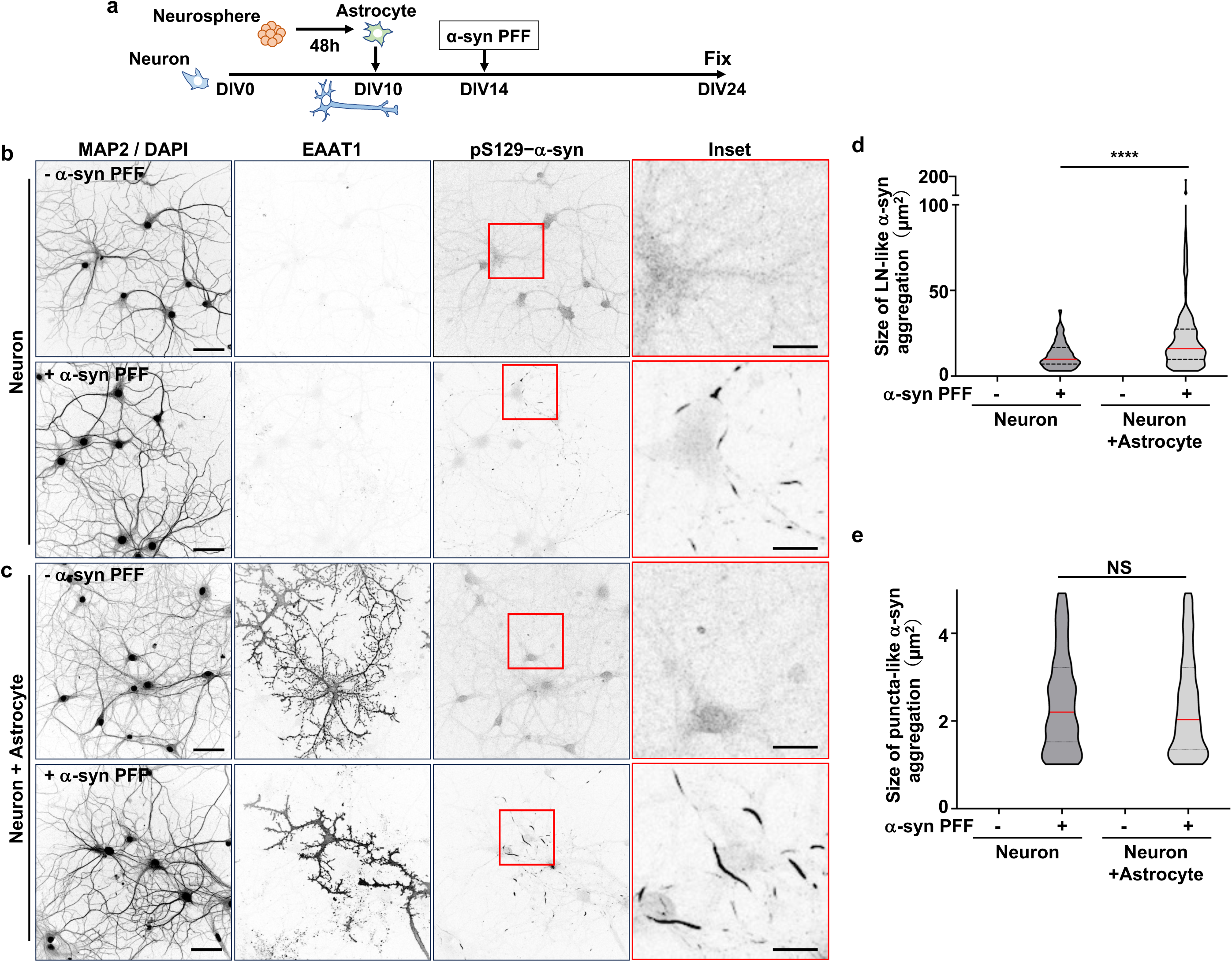
Astrocytes promote the elongation of LN-like α-syn aggregates in neurons. **(a)** Experimental design. Primary hippocampal neurons derived from embryonic mouse brains were cultured for 10 days in vitro (DIV10) and then co-cultured with astrocytes differentiated from neurospheres for an additional 14 days (DIV24). Co-cultures were treated at DIV14 with or without α-syn PFFs (0.5 μg/ml). **(b,c)** Effect of astrocytes on α-syn aggregate formation in neurons. After 10 days incubation with or without α-syn PFFs, neuron-only cultures (**b**) and neuron–astrocyte co-cultures (**c**) were fixed and immunostained for MAP2 (neurons), EAAT1 (astrocytes), and pS129–α-syn (aggregates). DAPI marks nuclei. Insets show higher magnification of boxed regions. Scale bar, 50 μm (MAP2/DAPI) and 20 μm (insets). **(d,e)** Quantification of LN-like (**d**) and puncta-like (**e**) α-syn aggregates. pS129–α-syn signal size was measured per field of view (n = 10 fields). Data are representative of three independent experiments. Red bars in violin plots indicate medians. Statistical significance was determined by Mann-Whitney test. ****p < 0.0001; NS, not significant.

To assess the specific contribution of astrocytes, we treated neuronal cultures and neuron–astrocyte co-cultures with or without α-syn PFFs at DIV14 and cultured them for an additional 10 days (DIV24) (Fig. 1a). Neuron-only cultures developed numerous pS129–α-syn-positive puncta and a limited number of short LN-like aggregates (Fig. 1b). In contrast, co-cultures with astrocytes exhibited a marked increase in both the length and abundance of LN-like structures (Fig. 1c). For quantitative analysis, we classified aggregates by size and shape: puncta were defined as structures with 1–5 μm^2^ in area, and LN-like aggregates as rod-shaped structures ≥5 μm^2^ in area with an ellipticity of ≤1/3. Most pS129–α-syn signals fell within these categories. Astrocyte co-culture significantly shifted the distribution of α-syn aggregates toward larger LN-like structures compared to neuron-only cultures (Fig. 1d), without altering the size distribution of puncta (Fig. 1e). These results suggest that astrocytes selectively promote the elongation and maturation of α-syn aggregates into LN-like forms within neurons.

### Astrocytes internalize and transfer α-syn PFFs to neurons

We next investigated whether astrocytes contribute to neuronal α-syn aggregation by internalizing and transferring exogenous α-syn seeds. Astrocytes were first pre-treated with α-syn PFFs for 24 hours and subsequently co-cultured with naïve, untreated neurons (Fig. 2a). Neurons co-cultured with untreated astrocytes showed no detectable pS129–α-syn signal (Fig. 2b). In contrast, neurons co-cultured with α-syn PFF-preloaded astrocytes developed pS129–α-syn-positive aggregates along their neurites after 14 days (Fig. 2c). The extent of aggregation increased over time, consistent with progressive seeding and propagation process (Fig. 2d). These results suggest that astrocytes, following α-syn PFF uptake, can transmit seeding-competent α-syn species to neighboring neurons, thereby initiating aggregate formation.

**Fig. 2.**
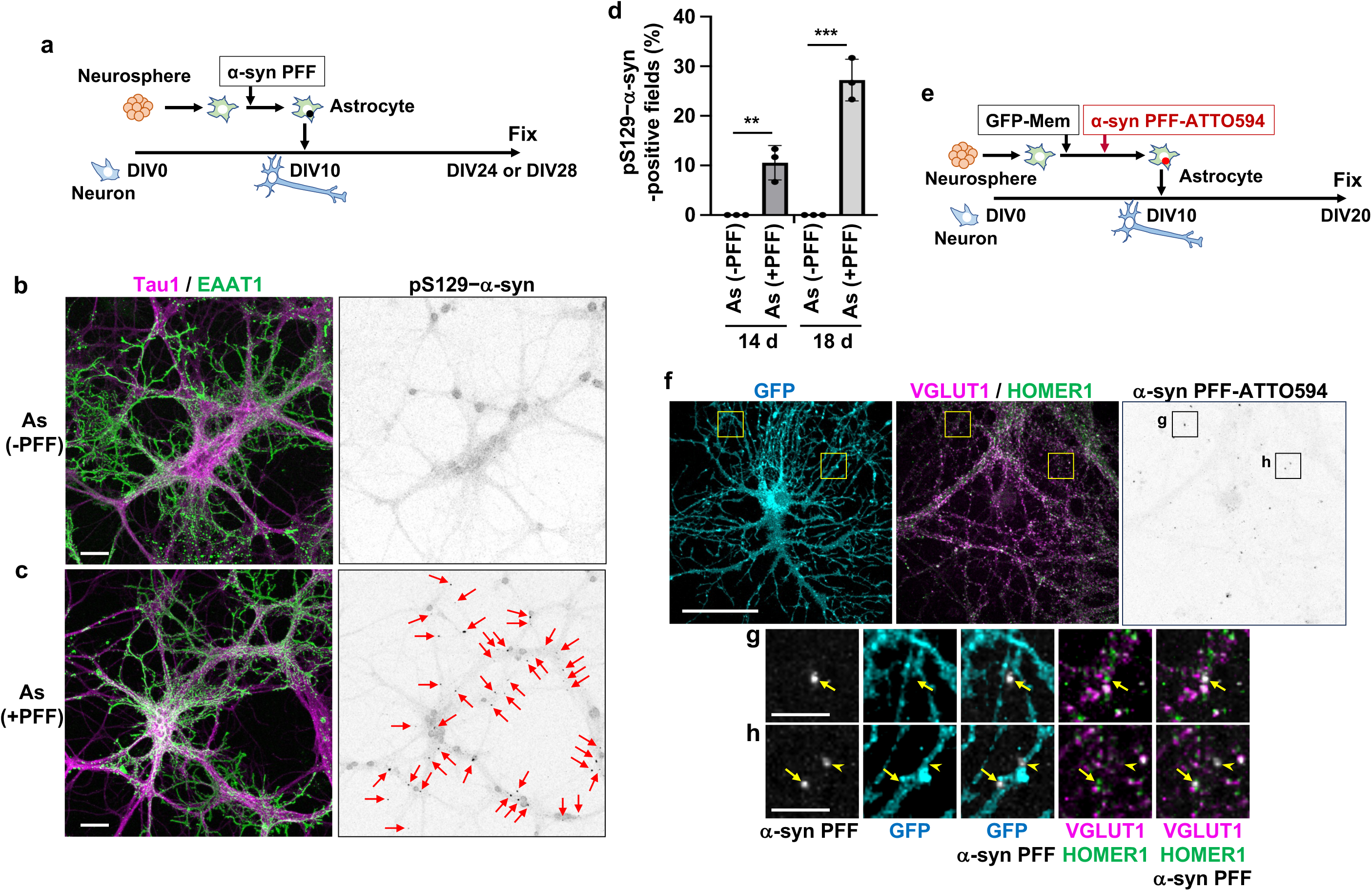
Astrocytes internalize and transfer pathogenic α-syn PFFs to neurons. **(a)** Experimental design. Astrocytes differentiated from neurospheres were treated with or without α-syn PFFs (0.5 μg/ml) for 24 h, passaged, and co-cultured with naïve neurons from DIV10 to DIV24 or DIV28. **(b,c)** Effect of α-syn PFF-preloaded astrocytes on α-syn aggregate formation in neurons. Neurons were co-cultured with untreated (**b**) or PFF-preloaded (**c**) astrocytes, fixed at DIV24, and immunostained for Tau1 (neurons, magenta), EAAT1 (astrocytes, green), and pS129–α-syn (aggregates, inverted grayscale). As (-PFF) represents untreated astrocytes, and As (+PFF) represents α-syn PFF-preloaded astrocytes. Arrows indicate α-syn aggregates in neurons. Scale bar, 50 μm. **(d)** Quantification of α-syn aggregate formation in neurons. The percentage of pS129–α-syn-positive fields was calculated (n = 60 fields per experiment). Data are combined from three independent experiments and analyzed by Mann-Whitney test. Error bars represent s.d. **p < 0.01, ***p < 0.001. **(e)** Experimental design for α-syn PFF transfer assay. Astrocytes were transfected with GFP-Mem, and 24 h later, they were treated with or without ATTO594-labeled α-syn PFFs (0.5 μg/ml) for 24 h, passaged, co-cultured with naïve neurons at DIV10, and maintained until DIV20. **(f-h)** Transfer of α-syn PFFs from astrocytes to neurons. Neurons co-cultured with α-syn PFF-ATTO594-preloaded astrocytes were fixed at DIV20 and immunostained for VGLUT1 (presynapse, magenta) and HOMER1 (postsynapse, green). GFP marks astrocytes. Magnified views of boxed regions are shown in **g** and **h**. Arrows indicate α-syn PFF puncta near synaptic markers, and arrowheads indicate puncta on astrocytic processes. Scale bars, 50 μm (**f**) and 10 μm (**g,h**).

To directly visualize this intercellular transfer, we labeled astrocytic membranes by expressing a membrane-targeted GAP43 fragment fused to green fluorescent protein (GFP-Mem)^48^ and treated these astrocytes with ATTO594-conjugated α-syn PFFs for 24 hours. After removal of unbound PFFs, the astrocytes were co-cultured with neurons (Fig. 2e). Fluorescence microscopy revealed ATTO594-positive puncta within GFP-labeled astrocytes (Fig. S3, arrowheads), as well as on Tau-positive neuronal axons (Fig. S3, arrows), indicating successful transfer of α-syn PFFs. No such puncta were detected in control co-cultures with untreated astrocytes (Fig. S3). Notably, a significant proportion of transferred α-syn puncta localized near synaptic markers, including VGLUT1 and HOMER1 (Fig. 2f–h, arrows), suggesting that tripartite synaptic contacts may serve as sites for astrocyte-to-neuron α-syn transfer. Together, these results provide direct evidence that astrocytes can internalize exogenous α-syn PFFs and relay them to neurons, thereby contributing to the initiation and propagation of α-syn pathology in a non-cell-autonomous manner.

### Astrocytic Cathepsin D cleaves the C-terminus of α-syn PFFs

C-terminally truncated α-syn species are enriched in astrocytes in PD brains^20,21^ and have been shown to possess enhanced seeding and aggregation potential^39,40,44,46^. We hypothesized that astrocytes actively generate these pathogenic fragments by cleaving the C-terminus of internalized α-syn PFFs. To test this, we treated astrocytes derived from neurospheres with α-syn PFFs and analyzed cellular lysates by immunoblotting at various time points. Using an antibody against the central region of α-syn, we detected full-length α-syn along with smaller molecular weight fragments in both the soluble and insoluble fractions as early as 6 hours after PFF exposure (Fig. 3a, arrow and arrowheads). These lower bands were not detected using an antibody specific for the C-terminal epitope, unequivocally confirming that they represent C-terminally truncated α-syn species. The abundance of truncated fragments increased over time, coinciding with a gradual loss of full-length protein (Fig. 3a), consistent with progressive proteolytic cleavage within astrocytes. These results indicate that astrocytes efficiently generate C-terminally truncated α-syn species upon internalization of PFFs.

**Fig. 3.**
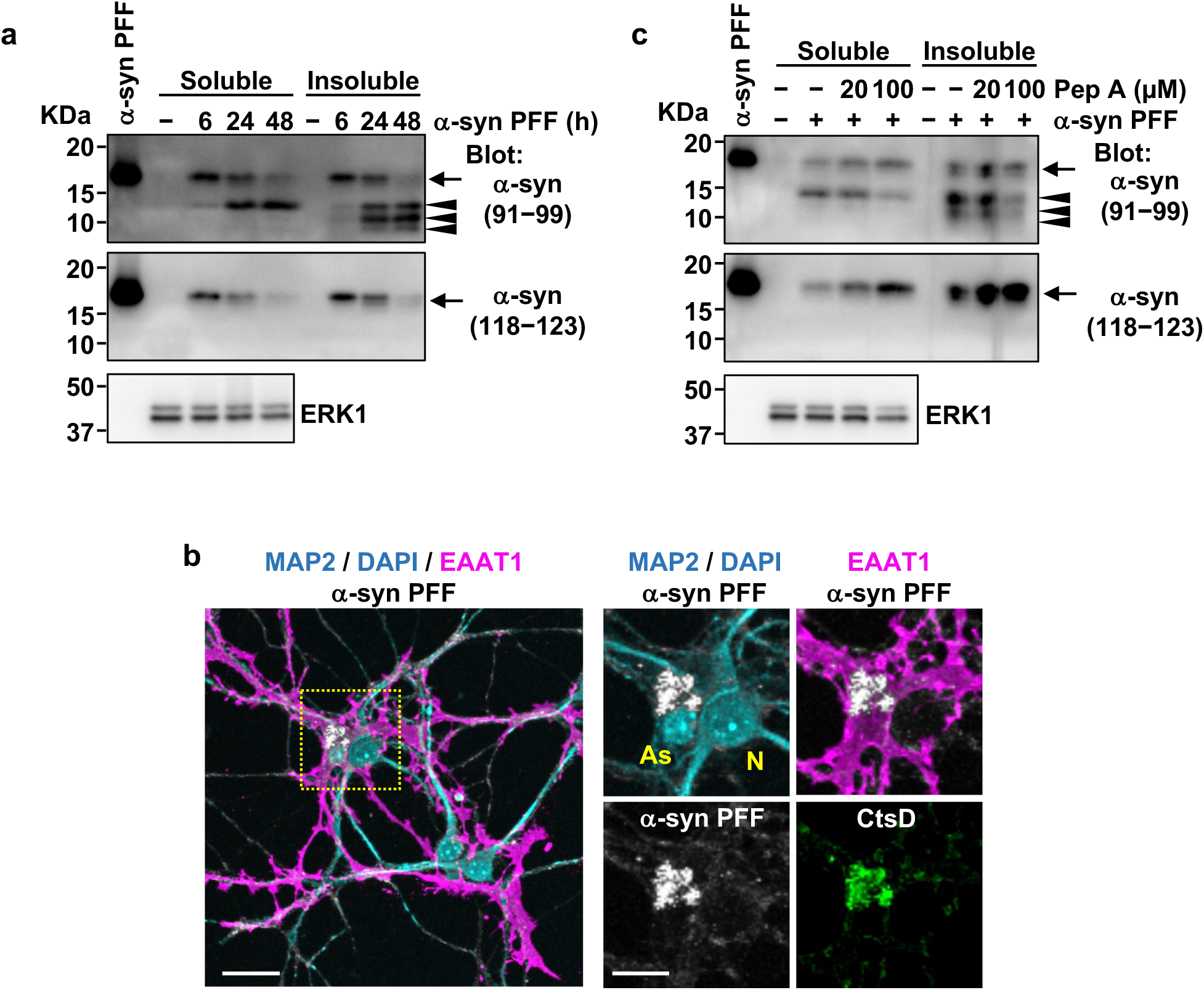
Astrocytic Cathepsin D cleaves the C-terminus of α-syn PFFs. **(a)** C-terminal cleavage of α-syn PFFs in astrocytes. Astrocytes differentiated from neurospheres were treated with α-syn PFFs (0.5 μg/ml) for the indicated times. Soluble and insoluble fractions from cell lysates were immunoblotted with antibodies recognizing the central region (amino acids 91–99) or the C-terminus (amino acids 118–123) of α-syn. Arrows indicate full-length α-syn and arrowheads indicate C-terminally truncated fragments. ERK serves as a loading control. **(b)** Co-localization of internalized α-syn PFF-ATTO594 with CtsD in astrocytes. Neuron–astrocyte co-cultures were incubated with α-syn PFF-ATTO594 (0.5 μg/ml) for 10 days, fixed at DIV24, and immunostained for MAP2 (neurons, cyan), EAAT1 (astrocytes, magenta), and CtsD (green). Higher-magnification views of the boxed region are shown at right. DAPI (cyan) marks nuclei. As, astrocyte nucleus; N, neuronal nucleus. Scale bar, 20 μm (left) and 10 μm (right). **(c)** Effect of Pep A on C-terminal cleavage of α-syn PFFs in astrocytes. Astrocytes were treated with α-syn PFFs (0.5 μg/ml) for 24 h in the presence or absence of Pep A (20 or 100 μM), and cell lysates were immunoblotted as in **a**.

We next sought to identify the protease responsible for this truncation. α-syn PFFs localize to lysosomes, where C-terminal cleavage is known to occur^44,46^, and we confirmed that astrocyte-internalized α-syn PFFs localized to lysosomes (Fig. S4). The lysosomal aspartic protease CtsD has previously been implicated in α-syn processing^35^. Immunofluorescence analysis revealed strong co-localization of internalized α-syn PFFs with CtsD in astrocytes (Fig. 3b, inset). To assess the functional role of CtsD, we treated PFF-exposed astrocytes with pepstatin A (Pep A), a selective CtsD inhibitor. Pep A treatment resulted in a dose-dependent reduction in the levels of C-terminally truncated α-syn fragments and preservation of full-length α-syn (Fig. 3c). These results indicate that CtsD is the principal enzyme responsible for C-terminal cleavage of α-syn PFFs in astrocytes.

### Astrocytic CtsD activity is required for the promotion of LN-like aggregation in neurons

We next examined whether CtsD activity in astrocytes is functionally required for their ability to promote α-syn aggregation in neurons. To this end, α-syn PFFs were added to neuron-only cultures or neuron–astrocyte co-cultures in the presence or absence of Pep A (Fig. 4a). In neuron-only cultures, inhibition of CtsD had no significant effect on the formation of either puncta or LN-like aggregates (Fig. 4b,c; Fig. S5), indicating that neuronal CtsD does not substantially contribute to PFF-induced aggregation under these conditions. By contrast, in co-cultures, Pep A treatment markedly reduced the size of LN-like aggregates to levels observed in neuron-only conditions (Fig. 4b,d; Fig. S5), while the formation of puncta remained unchanged (Fig. 4c,d). These results definitely demonstrate that CtsD activity specifically within astrocytes is essential for their ability to promote the elongation of LN-like α-syn aggregates in neurons. This suggests a model in which astrocytic proteolysis of PFFs generates aggregation-prone α-syn species that are subsequently transferred to neurons to potentiate pathology.

**Fig. 4.**
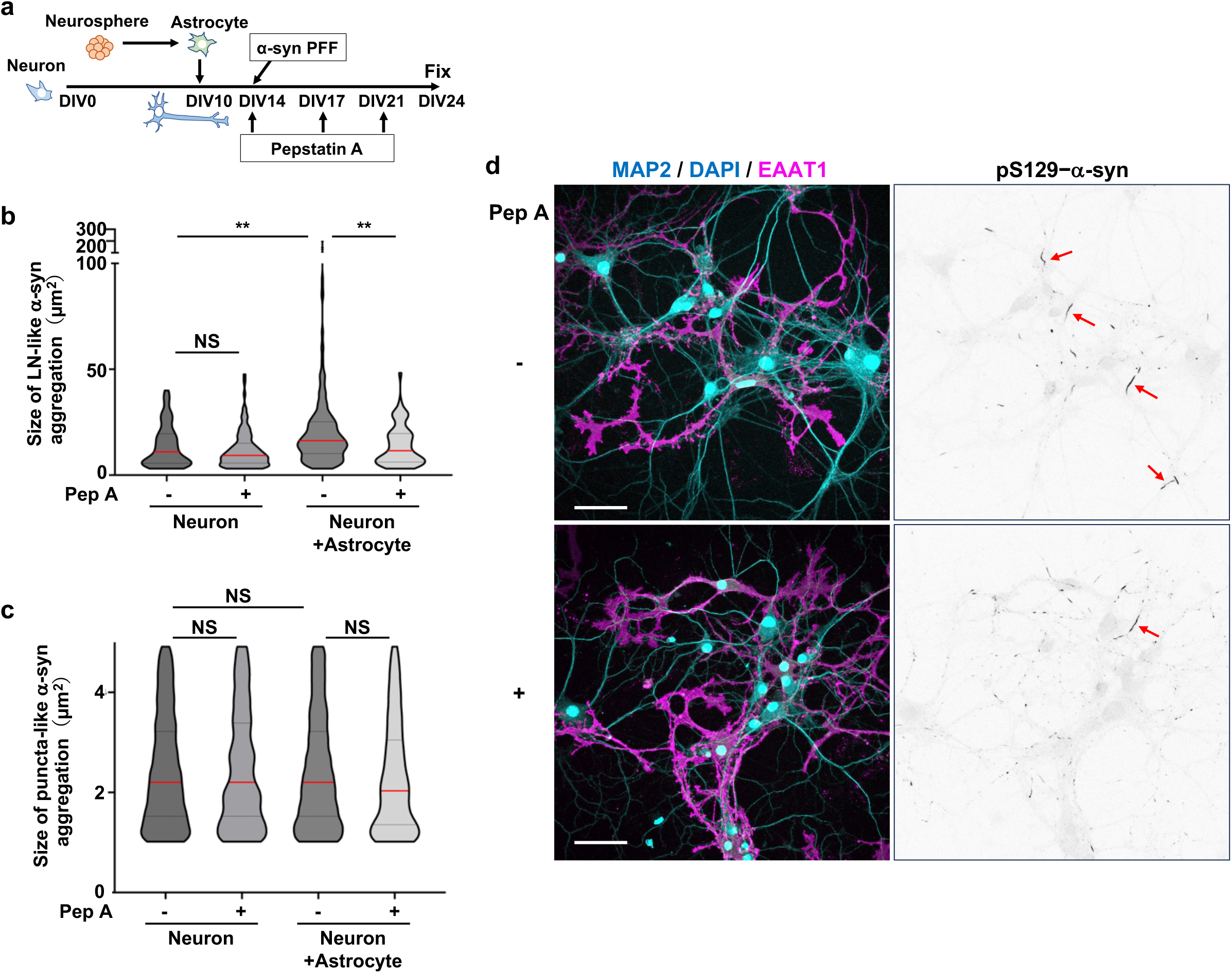
Cathepsin D activity is required for astrocyte-mediated promotion of LN-like aggregates in neurons. **(a)** Experimental design. Neuron–astrocyte co-cultures were treated with α-syn PFFs (0.5 μg/ml) for 10 days in the presence or absence of Pep A (10 μM), replenished every 3–4 days. **(b,c)** Quantification of LN-like (**b**) and puncta-like (**c**) α-syn aggregates. pS129–α-syn signal size was measured per field of view (n = 10 fields). Data are representative of three independent experiments. Red bars in violin plots indicate medians. Statistical significance was determined by Kruskal-Wallis test and Dunn’s multiple-comparison test. **p < 0.01; NS, not significant. **(d)** Effect of Pep A on astrocyte-mediated α-syn aggregation in neurons. Co-cultures were treated with α-syn PFFs in the presence or absence of Pep A (10 μM) and immunostained for MAP2 (neurons, cyan), EAAT1 (astrocytes, magenta), and pS129–α-syn (aggregates, inverted grayscale). DAPI (cyan) marks nuclei. Arrows indicate large LN-like α-syn aggregates (>30 μm^2^) in neurons. Scale bar, 50 μm.

### α-syn PFF uptake induces a senescence-like state and upregulation of CtsD in astrocytes

Astrocytes that internalized α-syn PFFs displayed marked morphological changes, including retraction of cellular processes and hypertrophy of the soma (Fig. 1c), features reminiscent of cellular senescence^49,50^. This observation is consistent with recent studies showing that fibrillar protein aggregates such as α-syn and Tau PFFs can trigger senescence-like phenotypes in glial cells^51,52^. To further characterize this phenotype, we assessed canonical markers of cellular senescence, including senescence-associated β-galactosidase (SA-β-gal) activity and intracellular reactive oxygen species (ROS) levels^53,54^. Astrocytes exposed to α-syn PFFs for 4 days exhibited a robust increase in SA-β-gal activity compared to untreated controls (Fig. 5a,b). In parallel, PFF treatment led to a significant elevation of ROS accumulation (Fig. 5a,c). These results support the conclusion that α-syn PFFs induce a senescence-like state in astrocytes.

**Fig. 5.**
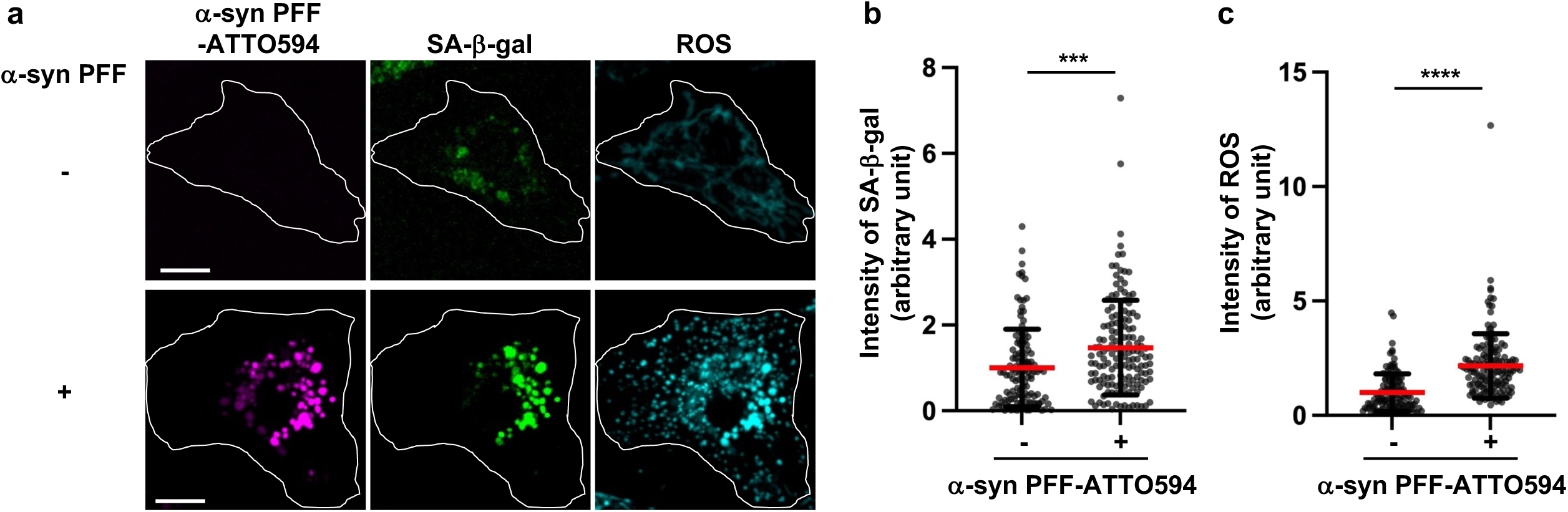
Uptake of α-syn PFFs induces a senescence-like phenotype in astrocytes. **(a)** Effect of α-syn PFFs on SA-β-gal activity and ROS levels. Astrocytes differentiated from neurospheres were treated with or without α-syn PFF-ATTO594 (0.5 μg/ml) for 4 days and stained for SPiDER-β-gal (SA-β-gal activity, green) and CellROX Deep Red (ROS levels, cyan). The boundaries of the astrocytes are indicated by white lines. Scale bar, 10 μm. **(b,c)** Quantification of SA-β-gal activity (**b**) and total cellular ROS levels (**c**). Data are mean ± s.d. (n = 125 cells) and representative of three independent experiments. Statistical significance was determined by Mann-Whitney test. ***p < 0.001, ****p < 0.0001.

Lysosomal dysfunction is a known driver of cellular senescence^55^, and α-syn PFFs have been reported to disrupt lysosomal integrity in HeLa cells or SH-SY5Y neuroblastoma cells^56,57^. We therefore examined whether the senescent phenotype observed in astrocytes might result from lysosomal damage.

Immunostaining for galectin-3 (Gal-3), a β-galactoside-binding lectin that binds glycans on the luminal side of lysosomal membrane proteins, labels lysosomes with ruptured membranes (Fig. 6a) ^58^. As early as 1 day after α-syn PFF treatment, Gal-3-positive lysosomes containing α-syn PFFs were detected, indicating membrane rupture (Fig. 6b). By day 6, Gal-3 puncta co-localizing with α-syn PFFs became enlarged and clustered in the perinuclear region (Fig. 6b), consistent with progressive lysosomal stress. Strikingly, these damaged lysosomes exhibited increased immunoreactivity for CtsD (Fig. 6b; Fig. S6a). Immunoblotting confirmed a rapid upregulation of CtsD protein within 24 hours of α-syn PFF exposure (Fig. 6c; Fig. S6b). These results implicate lysosomal membrane rupture as a trigger for CtsD upregulation in astrocytes, potentially forming a pathogenic feedback loop that enhances α-syn processing and propagation.

**Fig. 6.**
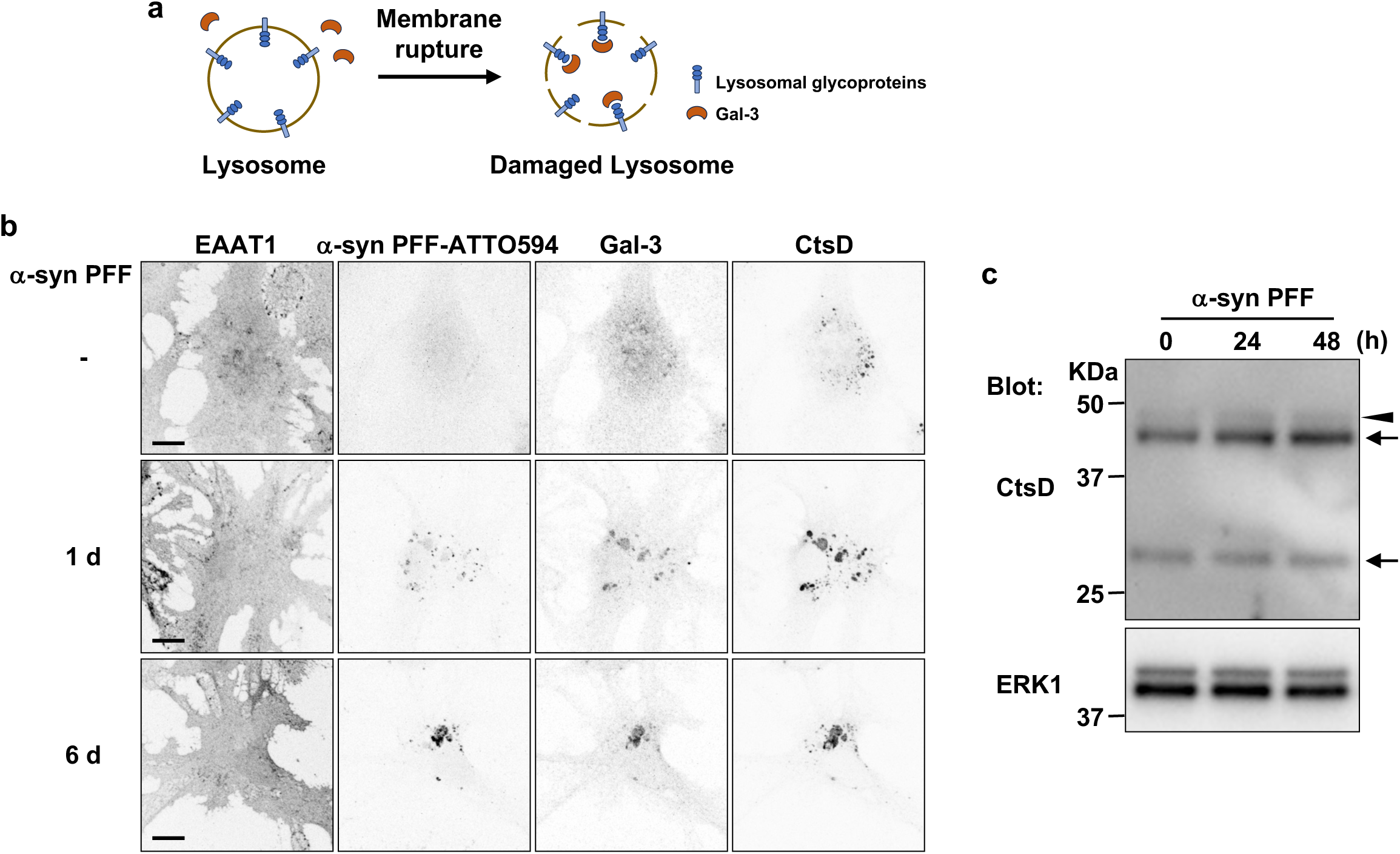
Uptake of α-syn PFFs disrupts lysosomal membrane integrity and upregulates CtsD in astrocytes. **(a)** Schematic of galectin-3 (Gal-3) labeling of lysosomal proteins. Gal-3 is a β-galactoside-binding lectin that binds glycans on the luminal side of lysosomal membrane proteins. Intact lysosomes are not labeled (left). Upon lysosomal membrane rupture, luminal glycans are exposed and Gal-3 binds (right). **(b)** Effect of α-syn PFFs on lysosomal integrity and CtsD expression. Astrocytes differentiated from neurospheres were treated with α-syn PFF-ATTO594 (0.5 μg/ml) for the indicated days and immunostained for EAAT1 (astrocytes), Gal-3 (lysosomal membrane rupture), and CtsD. Scale bar, 10 μm. **(c)** Effect of α-syn PFFs on CtsD protein levels. Astrocytes were treated with α-syn PFFs (0.5 μg/ml) for the indicated times, and cell lysates were immunoblotted for CtsD. Arrows indicate the active forms (48 kDa or 33 kDa) of CtsD and arrowhead indicates pro-CtsD. ERK serves as a loading control.

## DISCUSSION

Our study reveals a previously underappreciated, active role for astrocytes in promoting α-syn pathology. Using a physiologically relevant primary neuron– astrocyte co-culture system, we demonstrate that astrocytes not only internalize α-syn PFFs but also enzymatically process them via CtsD-mediated C-terminal truncation. These truncated, aggregation-prone α-syn species are subsequently transferred to neighboring neurons, where they accelerate the formation of LN-like inclusions. Furthermore, α-syn PFF exposure induces a senescence-like state and CtsD upregulation in astrocytes, suggesting a maladaptive feedback loop whereby lysosomal stress exacerbates the generation and spread of pathogenic α-syn. Together, these findings reposition astrocytes as active amplifiers of α-syn pathology, rather than passive bystanders or mere scavengers. A central strength of our study is the use of a co-culture model that recapitulates essential features of the in vivo neuron–glia microenvironment, including astrocytic morphological complexity and the formation of tripartite synapses. This system allowed us to dissect not only the intracellular handling of α-syn fibrils by astrocytes but also the functional consequences of astrocyte-to-neuron transfer of pathogenic species. We outline a sequential cascade: astrocytic uptake of α-syn PFFs, CtsD-mediated truncation into seeding-competent fragments, and intercellular transmission that fuels neuronal aggregate elongation.

Previous work has primarily characterized CtsD as a neuroprotective lysosomal protease, essential for α-syn degradation and overall proteostasis^34–38^. Indeed, CtsD deficiency in mice leads to spontaneous α-syn accumulation and neurodegeneration^34,36,38^. However, our data add critical nuance to this view by demonstrating that under pathological conditions, specifically, following the uptake of α-syn fibrils, CtsD cleaves the C-terminus of α-syn, generating fragments with heightened aggregation potential. These truncated species are enriched in both neurons and glia in PD and DLB brains^2,20,21,43^. Importantly, CtsD appears to fully degrade soluble α-syn under basal conditions but acts as a selective C-terminal protease in the context of fibrillar substrates^41^. This context-dependent switch suggests a dual role for CtsD: protective under physiological conditions, yet pathogenic under pathological conditions where α-syn fibrils are present. A key mechanistic insight from our study is that astrocyte-mediated promotion of α-syn aggregation is not merely a function of uptake but is specifically driven by CtsD-dependent proteolysis. Inhibition of CtsD selectively suppressed the elongation of LN-like aggregates in neurons without affecting punctate forms, highlighting a role for astrocytic enzymatic modification in shaping the morphology and progression of α-syn pathology. It is plausible that C-terminally truncated α-syn species serve as more efficient seeds for fibril growth, or that they resist neuronal clearance mechanisms, such as autophagy, thereby accumulating and promoting aggregation^59^.

Our findings also intersect with growing interest in glial senescence in neurodegenerative disease. Aging is a major risk factor for PD, and senescent astrocytes have been reported in human PD brains^60^. We show that α-syn PFF exposure triggers hallmarks of senescence in astrocytes, including increased SA-β-gal activity and ROS accumulation. These changes coincide with lysosomal membrane permeabilization and upregulation of CtsD, further implicating lysosomal dysfunction as a driver of both astrocytic aging and pathological α-syn processing. Given that senescent astrocytes are known to adopt a pro-inflammatory secretory phenotype and reduce neurotrophic support, their emergence in response to α-syn burden may contribute to a deleterious microenvironment that accelerates disease progression^61^.

In summary, our findings redefine the role of astrocytes in PD pathogenesis. Rather than acting solely as protective scavenger, astrocytes emerge as central players in α-syn propagation, capable of both enzymatic transformation and intercellular transmission of pathogenic species. The identification of CtsD as a molecular switch that converts astrocytic α-syn processing from protective to pathogenic underscores the importance of cellular context and lysosomal health in determining disease trajectory. While our model offers mechanistic clarity, future in vivo studies are essential to validate the pathological relevance of this astrocyte-driven axis and assess its therapeutic tractability. Notably, the link between astrocytic senescence and α-syn propagation raises the possibility that targeting glial aging or lysosomal resilience may represent novel strategies to interrupt disease progression in PD and related synucleinopathies.

## Methods

### Antibodies and reagents

The antibodies and suppliers were as follows: anti-MAP2 (ab5392, abcam); anti-EAAT1 (D44E2, CST; GLAST-GP-Af1000, Frontier Institute); anti-Tau1 (Merck Millipore); anti-VGLUT1 (Synaptic Systems); anti-HOMER1 (Synaptic Systems); anti-α-syn (Syn1, BD Transduction; MJFR1, Abcam); anti-pS129–α-syn (EP1536Y, Abcam); anti-CtsD (R&D Systems); anti-Lamp1 (1D4B, DSHB); anti-Gal-3 (sc-23938, Santa Cruz); anti-ERK1 (sc-94, Santa Cruz). Human α-syn PFF (SPR-322) and ATTO594-conjugated α-syn PFF (SPR-322-A594) were purchased from StressMarq Biosciences. Pepstatin A was purchased from Selleck Chemicals. pAcGFP-Mem was obtained from Takara Bio. HB-EGF was purchased from Peprotech. C57BL/6J mice were obtained from SLC.

### Cell cultures

Primary mouse hippocampal neurons were prepared as described previously^33^ and cultured in Neurobasal medium (Thermo Fisher) supplemented with B27 (Thermo Fisher), 2 mM GlutaMAX (Thermo Fisher), and gentamicin (Merck Millipore). Cytosine arabinoside (2 μM; Merck Millipore) was added at DIV2. Astrocytes were differentiated from neurospheres prepared from the ganglionic eminence of embryonic day 18.5 (E18.5) mice. For differentiation of astrocytes, neurosphere were passaged and cultured for 2–3 days in a medium composed of neuron medium and MEM (Thermo Fisher) (ratio 1:4) supplemented with 1% fetal bovine serum (FBS; Corning), as described previously^33^. Astrocytes were transfected with pAcGFP1-Mem using Lipofectamine 2000 (Thermo Fisher) according to the manufacturer’s protocol. In vitro co-culture system for neurons and astrocytes were prepared as described previously^33^. Briefly, hippocampal neurons (6 x 10^4^ cells/well) were seeded on poly-L-lysine (PLL)-coated cover glass in 24-well plates. Astrocytes (1 x 10^4^ cells/well) were added at DIV10 and co-cultured for the indicated periods.

### Immunofluorescence and image analysis

Cells grown on coverslips were fixed in 4% paraformaldehyde for 15 min at 37°C, permeabilized with 0.5% Triton X-100 for 5 min, and incubated with the indicated primary and fluorophore-conjugated secondary antibodies. Primary antibodies included: chicken anti-MAP2 (1:2000), guinea pig anti-EAAT1 (1:1000), anti-VGLUT1 (1:1000), rabbit anti-EAAT1 (1:300), anti-pS129–α-syn (1:500), mouse anti-Tau1 (1:1000), anti-HOMER1 (1:100), rat anti-Lamp1 (1:50), anti-Gal-3 (1:1000), and goat anti-CtsD (1:50). The secondary antibodies included DyLight 405 donkey anti-chicken IgG, donkey anti-guinea pig IgG, Alexa-Fluor 488-, 555-, or 647-conjugated antibodies against mouse, rabbit rat, or goat IgG (Thermo Fisher). Confocal images were acquired using an Olympus FV3000 microscope with either 20x NA 0.75 objective (1.06 μm intervals) or 60x oil-immersion NA 1.4 objective (0.42 μm intervals). Z-stacks were processed using FV3000 software. Imaging parameters were optimized for the brightest unsaturated signal and held constant for all conditions within an experiment. For quantification of CtsD expression in astrocytes, regions of interest (ROIs) were drawn for individual cells, and mean fluorescence intensity of CtsD was measured using Fiji/Image J (NIH). Experiments were repeated at least three times.

### Quantification of α-syn aggregates

pS129–α-syn fluorescence was quantified using the Fiji/ImageJ plugin (NIH). Z-stack images were background-subtracted (rolling ball radius) and binarized (MaxEntropy threshold). Fitted ellipses were used to measure aggregate area and ellipticity. Aggregates of 1–5 μm^2^ were defined as “puncta”, and rod-shaped aggregates ≥5 μm^2^ with ellipticity of ≤1/3 were classified as “LN-like structures”. At least 10 fields were analyzed per experiment, with a minimum of three independent replicates.

### SA-β-galactosidase activity and ROS measurement

SA-β-gal activity was assessed using the SPiDER-β-gal fluorometric kit (Dojindo), and ROS levels were measured with CellRox Deep Red (Thermo Fisher), following the manufacturers’ instructions. Briefly, astrocytes were seeded in μ-Slide 8-well chambers (ibidi) and treated with or without α-syn PFF-ATTO594 for 4 days in neuron medium/MEM (1:4) supplemented with N2 (Thermo Fisher), HB-EGF (5 ng/ml), and 0.1% FBS. Cells were incubated with bafilomycin A1 (1 h), followed by SPiDER-β-gal and CellROX staining (30 min). Images were acquired with an Andor Dragonfly spinning disk confocal microscope using a 60x oil-immersion NA 1.4 objective. Regions of interest (ROIs) were drawn for individual cells, and mean fluorescence intensity was quantified using Fiji/Image J (NIH). Experiments were repeated at least three times.

### Western blotting analysis

Astrocytes were cultured in neuron medium/MEM (1:4) supplemented with N2, HB-EGF (5 ng/ml), and 0.1% FBS. Following α-syn PFF treatment, cells were lysed in RIPA buffer [50 mM Tris-HCl (pH 7.4), 0.15 M NaCl, 0.25% deoxycholic acid, 1% NP-40, 1 mM EDTA, 1 mM dithiothreitol, phosphatase inhibitor cocktail 2 (Sigma), and protease inhibitor cocktail (Sigma)], and centrifuged at 15,000 *g* for 12 min. The supernatant was collected as the soluble fraction; the pellet was resuspended in 2% sodium dodecyl sulfate (SDS)/tris-buffered saline (TBS) as the insoluble fraction. Samples were separated by 5-20% gradient SDS-polyacrylamide gels (e-PAGEL, ATTO) and transferred to 0.45 μm PVDF membranes (Amersham). The membranes were fixed with 4% (w/v) paraformaldehyde in TBS for 30 min to prevent detachment of α-syn from the blotted membranes. Primary antibodies: anti-α-syn (1:500, BD Transduction; 1:1000, Abcam), anti-ERK1 (1:1000), anti-CtsD (1:200).

### Statistical analysis

Data were analyzed using Mann-Whitney test, Kruskal-Wallis test, and Dunn’s multiple-comparison test (GraphPad Prism 10). Error bars indicate s.d. Exact n values are provided in figure legends. No statistical method was used to predetermine sample size.

## ACKNOWLEDGEMENTS

This research was supported by grants from the Ministry of Education, Culture, Sports, Science and Technology of Japan and JST (Moonshot R&D; Grant Number JPMJMS2024) (to H.H.) and Japan Agency for Medical Research and Development (Multidisciplinary Frontier Brain and Neuroscience Discoveries (Brain/MINDS 2.0); Grant Number JP24wm0625317) (to H.H.).

## AUTHOR CONTRIBUTIONS

H.H., N.H., T.K., M.M., K.M., Y.T. and K.M. designed the experiments and analyzed the data. H.H. and K.M. wrote the manuscript. H.H., K.O. and H.S. performed the experiments.

## SUPPLEMENTARY FIGURE LEGENDS

**Fig. S1.**
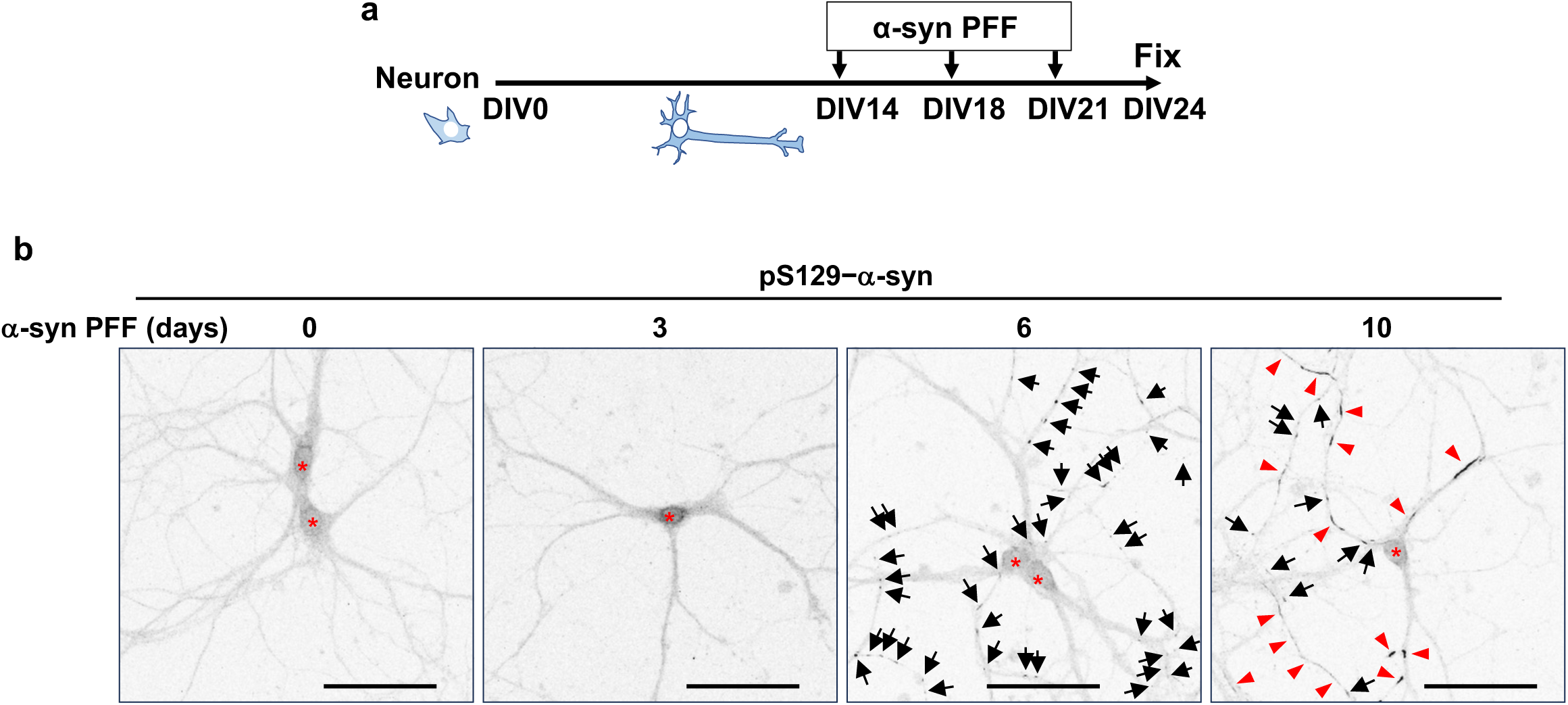
α-syn PFFs induce α-syn aggregate formation in neurons. **(a)** Experimental design. Primary hippocampal neurons were treated once with α-syn PFFs (0.5 μg/ml) at DIV14, DIV18, DIV21. All cultures were fixed at DIV24. **(b)** Aggregate formation in neurons following α-syn PFFs treatment. After treatment with α-syn PFFs for the indicated days, neurons were fixed and immunostained for MAP2 (neurons) and pS129–α-syn (aggregates, inverted grayscale). Arrows indicate punctate aggregates, arrowheads indicate LN-like aggregates, and asterisks indicate nuclei. Scale bar, 50 μm.

**Fig. S2.**
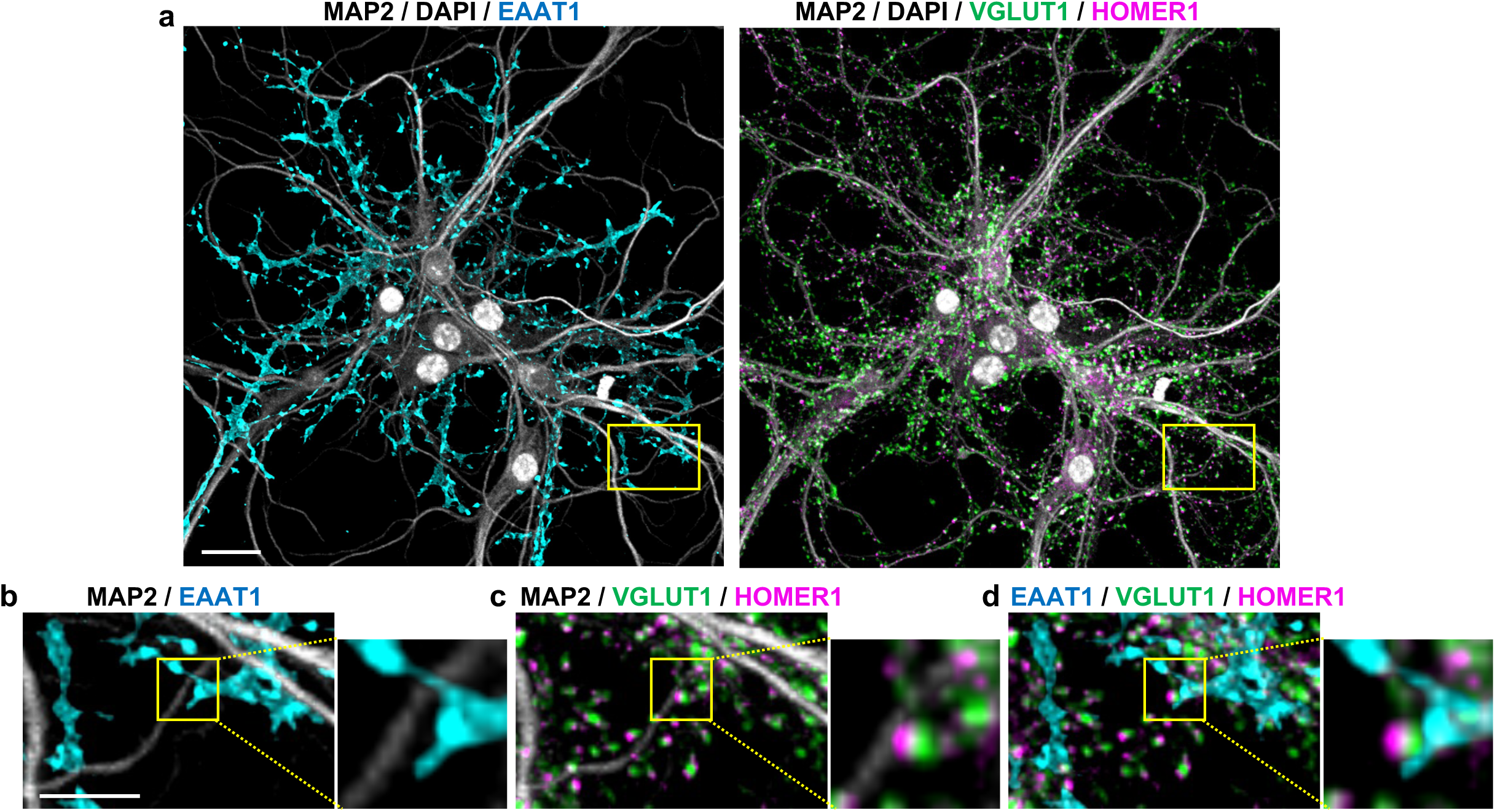
Characterization of primary mouse hippocampal neuron– astrocyte co-cultures. **(a)** Immunofluorescence images of neuron–astrocyte co-cultures. Primary hippocampal neurons derived from embryonic mouse brains were cultured for 10 days in vitro and then co-cultured with astrocytes differentiated from neurospheres for an additional 14 days. Cells were immunostained for MAP2 (neurons, gray), EAAT1 (astrocytes, cyan), VGLUT1 (presynapse, green), and HOMER1 (postsynapse, magenta). DAPI (gray) marks nuclei. Scale bar, 20 μm. **(b-d)** Magnified views of boxed regions in **a**. They show representative tripartite synapses. Scale bar, 10 μm.

**Fig. S3.**
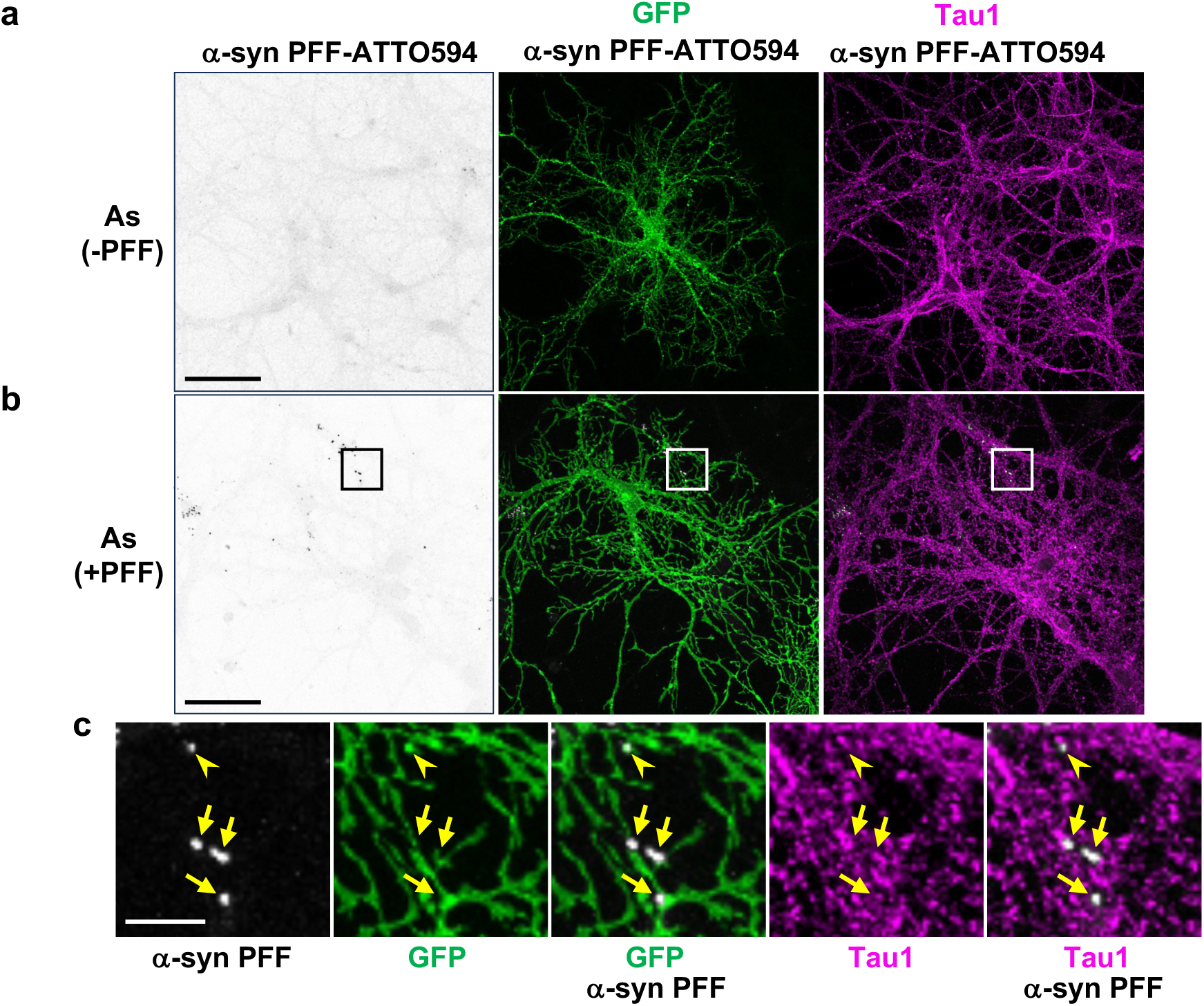
Astrocyte-to-neuron transfer of α-syn PFFs. **(a,b)** Transfer of α-syn PFFs from astrocytes to neurons. Astrocytes expressing GFP-Mem were treated with or without α-syn PFF-ATTO594 (0.5 μg/ml) for 24 h, passaged, and co-cultured with naïve neurons for 10 days. Neurons were fixed and immunostained for Tau1 (axon, magenta). GFP marks astrocytes. As (-PFF) represents untreated astrocytes, and As (+PFF) represents α-syn PFF-ATTO594-preloaded astrocytes. Scale bar, 50 μm. **(c)** Magnified views of boxed regions in **b**. Arrowheads indicate α-syn PFF-ATTO594 puncta on GFP-positive astrocytic processes, and arrows indicate puncta along neuronal axons. Scale bar, 10 μm.

**Fig. S4.**
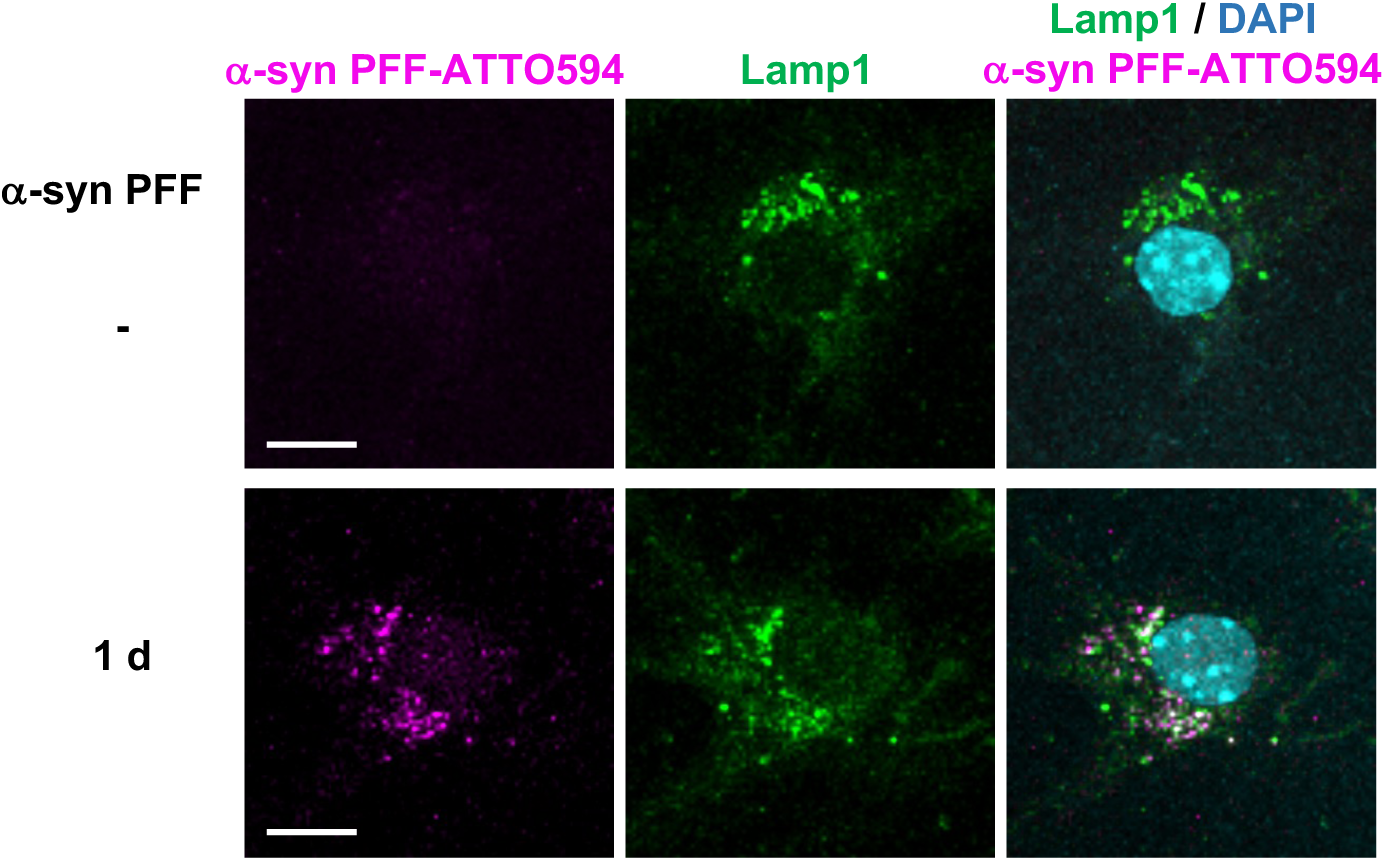
Localization of internalized α-syn PFFs in lysosomes. Astrocytes differentiated from neurospheres were treated with or without α-syn PFF-ATTO594 (0.5 μg/ml) for 1 day and immunostained for Lamp1 (lysosomes, green). DAPI (cyan) marks nuclei. Scale bar, 10 μm.

**Fig. S5.**
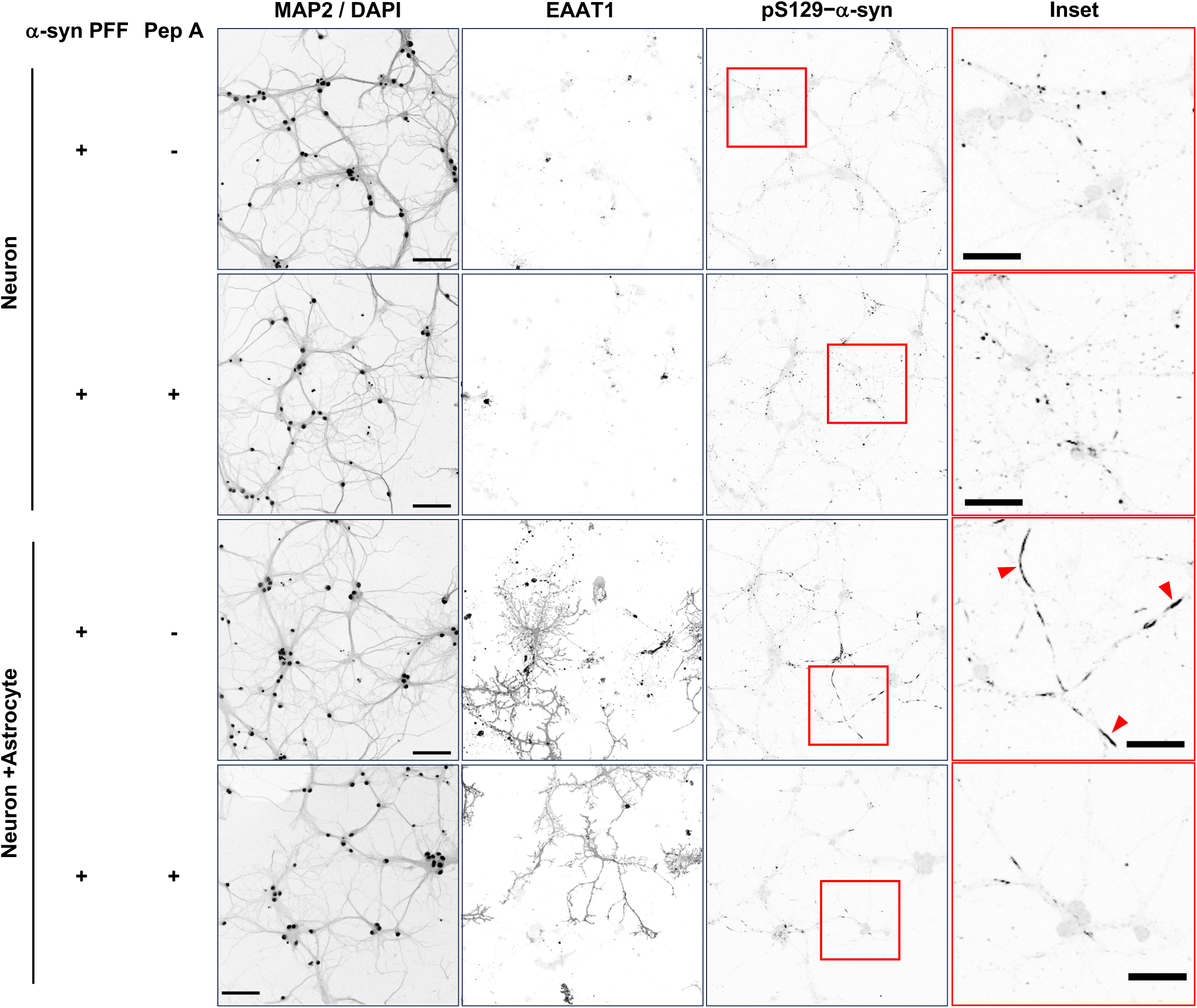
Effect of Pep A on neuronal α-syn aggregation. Neuron-only cultures and neuron–astrocyte co-cultures were treated with α-syn PFFs (0.5 μg/ml) in the presence or absence of Pep A (10 μM) for 10 days, replenished every 3–4 days. Cells were immunostained for MAP2 (neurons), EAAT1 (astrocytes), and pS129–α-syn (aggregates). DAPI marks nuclei. Insets show higher magnification of boxed regions. Arrowheads indicate large LN-like α-syn aggregates (>30 μm^2^) in neurons. Scale bar, 50 μm (left) and 25 μm (inset).

**Fig. S6.**
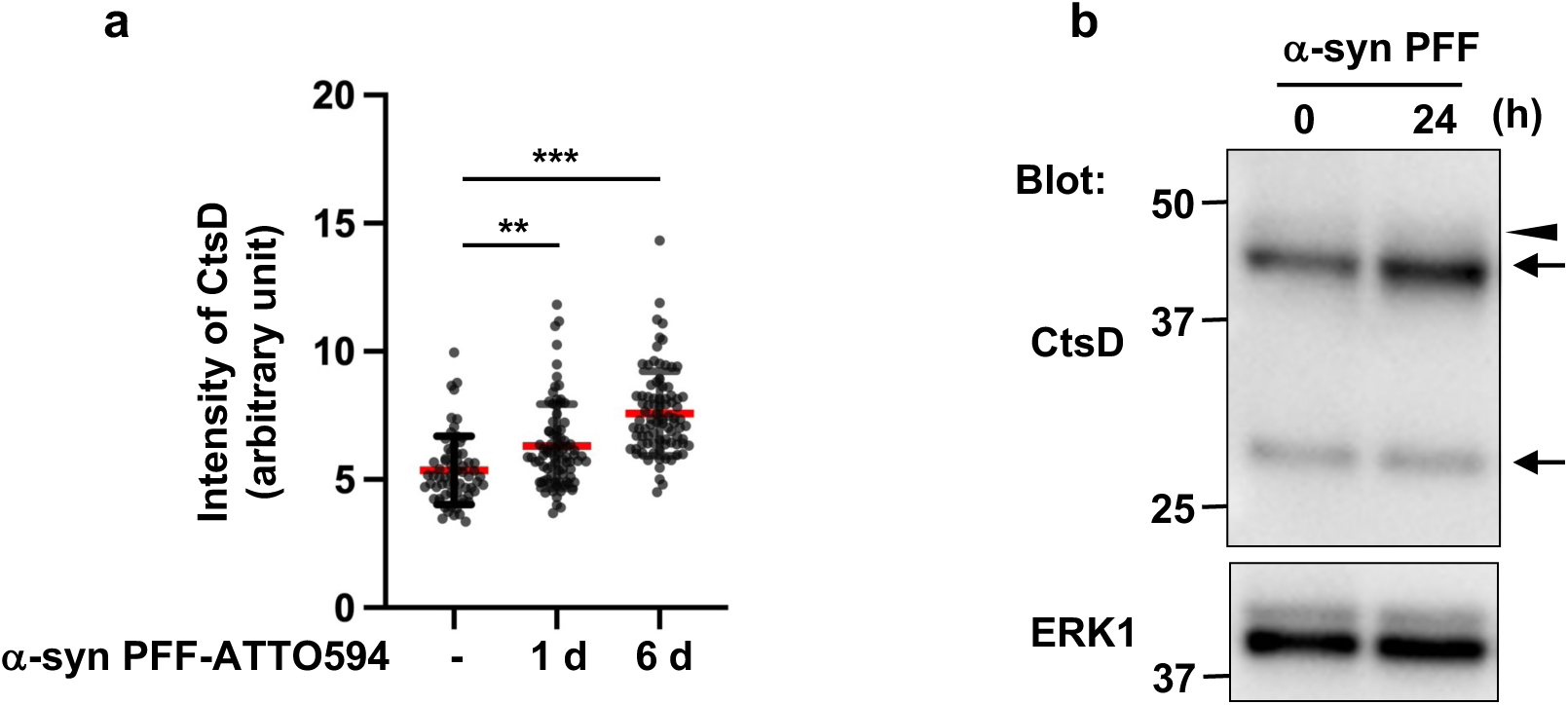
α-syn PFF uptake induces CtsD expression in astrocytes. **(a)** Quantification of CtsD expression. Astrocytes differentiated from neurospheres were treated with α-syn PFF-ATTO594 (0.5 μg/ml) for the indicated days and immunostained for CtsD. Data are mean ± s.d. (n = 80 cells) and representative of three independent experiments. Statistical significance was determined by Kruskal-Wallis test and Dunn’s multiple-comparison test. **p < 0.01, ***p < 0.001. **(b)** Effect of α-syn PFFs on CtsD protein levels. Astrocytes were treated with α-syn PFFs (0.5 μg/ml) for 24 h, and cell lysates were immunoblotted for CtsD. Arrows indicate the active forms (48 kDa or 33 kDa) of CtsD and arrowhead indicates pro-CtsD. ERK serves as a loading control.

## References

1. Spillantini, M. G. et al. α-Synuclein in Lewy bodies. Nature 388, 839–840 (1997).

2. Baba, M. et al. Aggregation of alpha-synuclein in Lewy bodies of sporadic Parkinson’s disease and dementia with Lewy bodies. Am. J. Pathol. 152, 879–884 (1998).

3. Braak, H., Sandmann-Keil, D., Gai, W. & Braak, E. Extensive axonal Lewy neurites in Parkinson’s disease: a novel pathological feature revealed by *α*-synuclein immunocytochemistry. Neurosci. Lett. 265, 67–69 (1999).

4. Braak, H. et al. Staging of brain pathology related to sporadic Parkinson’s disease. Neurobiol. Aging 24, 197–211 (2003).

5. Morris, H. R., Spillantini, M. G., Sue, C. M. & Williams-Gray, C. H. The pathogenesis of Parkinson’s disease. The Lancet 403, 293–304 (2024).

6. Maroteaux, L., Campanelli, J. T. & Scheller, R. H. Synuclein: a neuron-specific protein localized to the nucleus and presynaptic nerve terminal. J. Neurosci. 8, 2804–2815 (1988).

7. Abeliovich, A. et al. Mice lacking α-Synuclein display functional deficits in the nigrostriatal dopamine system. Neuron 25, 239–252 (2000).

8. Cabin, D. E. et al. Synaptic vesicle depletion correlates with attenuated synaptic responses to prolonged repetitive stimulation in mice lacking α-Synuclein. J. Neurosci. 22, 8797–8807 (2002).

9. Burré, J. et al. α-Synuclein promotes SNARE-complex assembly in vivo and in vitro. Science 329, 1663–1667 (2010).

10. Bendor, J. T., Logan, T. P. & Edwards, R. H. The function of α-Synuclein. Neuron 79, 1044–1066 (2013).

11. Burré, J., Sharma, M. & Südhof, T. C. Cell biology and pathophysiology of α-Synuclein. Cold Spring Harb. Perspect. Med. 8, a024091 (2018).

12. Li, J.-Y. et al. Lewy bodies in grafted neurons in subjects with Parkinson’s disease suggest host-to-graft disease propagation. Nat. Med. 14, 501–503 (2008).

13. Kordower, J. H., Chu, Y., Hauser, R. A., Freeman, T. B. & Olanow, C. W. Lewy body–like pathology in long-term embryonic nigral transplants in Parkinson’s disease. Nat. Med. 14, 504–506 (2008).

14. Luk, K. C. et al. Pathological α-Synuclein transmission initiates Parkinson-like neurodegeneration in nontransgenic mice. Science 338, 949–953 (2012).

15. Masuda-Suzukake, M. et al. Prion-like spreading of pathological α-synuclein in brain. Brain 136, 1128–1138 (2013).

16. Luk, K. C. et al. Exogenous α-synuclein fibrils seed the formation of Lewy body-like intracellular inclusions in cultured cells. Proc. Natl. Acad. Sci. 106, 20051–20056 (2009).

17. Volpicelli-Daley, L. A. et al. Exogenous α-Synuclein fibrils induce Lewy body pathology leading to synaptic dysfunction and neuron death. Neuron 72, 57–71 (2011).

18. Mahul-Mellier, A.-L. et al. The process of Lewy body formation, rather than simply α-synuclein fibrillization, is one of the major drivers of neurodegeneration. Proc. Natl. Acad. Sci. 117, 4971–4982 (2020).

19. Sofroniew, M. V. & Vinters, H. V. Astrocytes: biology and pathology. Acta Neuropathol. (Berl.) 119, 7–35 (2010).

20. Sorrentino, Z. A., Giasson, B. I. & Chakrabarty, P. α-Synuclein and astrocytes: tracing the pathways from homeostasis to neurodegeneration in Lewy body disease. Acta Neuropathol. (Berl.) 138, 1–21 (2019).

21. Altay, M. F., Liu, A. K. L., Holton, J. L., Parkkinen, L. & Lashuel, H. A. Prominent astrocytic alpha-synuclein pathology with unique post-translational modification signatures unveiled across Lewy body disorders. Acta Neuropathol. Commun. 10, 163 (2022).

22. Smethurst, P., Franklin, H., Clarke, B. E., Sidle, K. & Patani, R. The role of astrocytes in prion-like mechanisms of neurodegeneration. Brain 145, 17– 26 (2022).

23. Ozoran, H. & Srinivasan, R. Astrocytes and alpha-Synuclein: Friend or Foe? J. Park. Dis. 13, 1289–1301 (2023).

24. Araque, A., Parpura, V., Sanzgiri, R. P. & Haydon, P. G. Tripartite synapses: glia, the unacknowledged partner. Trends Neurosci. 22, 208–215 (1999).

25. Allen, N. J. & Eroglu, C. Cell biology of astrocyte-synapse interactions. Neuron 96, 697–708 (2017).

26. Loria, F. et al. α-Synuclein transfer between neurons and astrocytes indicates that astrocytes play a role in degradation rather than in spreading. Acta Neuropathol. (Berl.) 134, 789–808 (2017).

27. Tsunemi, T. et al. Astrocytes protect human dopaminergic neurons from α-Synuclein accumulation and propagation. J. Neurosci. 40, 8618–8628 (2020).

28. Lee, H.-J. et al. Direct Transfer of α-Synuclein from neuron to astroglia causes inflammatory responses in synucleinopathies. J. Biol. Chem. 285, 9262–9272 (2010).

29. Rannikko, E. H., Weber, S. S. & Kahle, P. J. Exogenous α-synuclein induces toll-like receptor 4 dependent inflammatory responses in astrocytes. BMC Neurosci. 16, 57 (2015).

30. Hughes, C. D. et al. Picomolar concentrations of oligomeric alpha-synuclein sensitizes TLR4 to play an initiating role in Parkinson’s disease pathogenesis. Acta Neuropathol. (Berl.) 137, 103–120 (2019).

31. Chou, T.-W. et al. Fibrillar α-synuclein induces neurotoxic astrocyte activation via RIP kinase signaling and NF-κB. Cell Death Dis. 12, 1–11 (2021).

32. Leandrou, E. et al. α-Synuclein oligomers potentiate neuroinflammatory NF-κB activity and induce Cav3.2 calcium signaling in astrocytes. Transl. Neurodegener. 13, 11 (2024).

33. Nozawa, O. et al. Necl2/3-mediated mechanism for tripartite synapse formation. Development 150, dev200931 (2023).

34. Qiao, L. et al. Lysosomal enzyme cathepsin D protects against alpha-synuclein aggregation and toxicity. Mol. Brain 1, 17 (2008).

35. Sevlever, D., Jiang, P. & Yen, S.-H. C. Cathepsin D Is the main lysosomal enzyme involved in the degradation of α-Synuclein and generation of its carboxy-terminally truncated species. Biochemistry 47, 9678–9687 (2008).

36. Cullen, V. et al. Cathepsin D expression level affects alpha-synuclein processing, aggregation, and toxicity in vivo. Mol. Brain 2, 5 (2009).

37. Vidoni, C., Follo, C., Savino, M., Melone, M. A. B. & Isidoro, C. The role of Cathepsin D in the pathogenesis of human neurodegenerative disorders. Med. Res. Rev. 36, 845–870 (2016).

38. Suzuki, C. et al. Lack of Cathepsin D in the central nervous system results in microglia and astrocyte activation and the accumulation of proteinopathy-related proteins. Sci. Rep. 12, 11662 (2022).

39. Murray, I. V. J. et al. Role of α-Synuclein carboxy-terminus on fibril formation in vitro. Biochemistry 42, 8530–8540 (2003).

40. Hoyer, W., Cherny, D., Subramaniam, V. & Jovin, T. M. Impact of the acidic C-terminal region comprising amino acids 109−140 on α-Synuclein aggregation in vitro. Biochemistry 43, 16233–16242 (2004).

41. McGlinchey, R. P. & Lee, J. C. Cysteine cathepsins are essential in lysosomal degradation of α-synuclein. Proc. Natl. Acad. Sci. 112, 9322–9327 (2015).

42. Campbell, B. C. V. et al. The solubility of α-synuclein in multiple system atrophy differs from that of dementia with Lewy bodies and Parkinson’s disease. J. Neurochem. 76, 87–96 (2001).

43. Li, W. et al. Aggregation promoting C-terminal truncation of α-synuclein is a normal cellular process and is enhanced by the familial Parkinson’s disease-linked mutations. Proc. Natl. Acad. Sci. 102, 2162–2167 (2005).

44. Sorrentino, Z. A. & Giasson, B. I. The emerging role of α-synuclein truncation in aggregation and disease. J. Biol. Chem. 295, 10224–10244 (2020).

45. Quintin, S. et al. Cellular processing of α-synuclein fibrils results in distinct physiological C-terminal truncations with a major cleavage site at residue Glu 114. J. Biol. Chem. 299, 104912 (2023).

46. Suthar, S. K. & Lee, S.-Y. Truncation or proteolysis of α-synuclein in Parkinsonism. Ageing Res. Rev. 90, 101978 (2023).

47. Volpicelli-Daley, L. A., Luk, K. C. & Lee, V. M.-Y. Addition of exogenous α-synuclein preformed fibrils to primary neuronal cultures to seed recruitment of endogenous α-synuclein to Lewy body and Lewy neurite–like aggregates. Nat. Protoc. 9, 2135–2146 (2014).

48. Moriyoshi, K., Richards, L. J., Akazawa, C., O’Leary, D. D. M. & Nakanishi, S. Labeling neural cells using adenoviral gene transfer of membrane-targeted GFP. Neuron 16, 255–260 (1996).

49. Mansour, H. et al. Aging-related changes in astrocytes in the rat retina: imbalance between cell proliferation and cell death reduces astrocyte availability. Aging Cell 7, 526–540 (2008).

50. Lawal, O., Ulloa Severino, F. P. & Eroglu, C. The role of astrocyte structural plasticity in regulating neural circuit function and behavior. Glia 70, 1467– 1483 (2022).

51. Verma, D. K. et al. Alpha-synuclein preformed fibrils induce cellular senescence in Parkinson’s disease models. Cells 10, 1694 (2021).

52. Ungerleider, K. et al. Astrocyte senescence and SASP in neurodegeneration: tau joins the loop. Cell Cycle 20, 752–764 (2021).

53. Dimri, G. P. et al. A biomarker that identifies senescent human cells in culture and in aging skin in vivo. Proc. Natl. Acad. Sci. 92, 9363–9367 (1995).

54. Muñoz-Espín, D. & Serrano, M. Cellular senescence: from physiology to pathology. Nat. Rev. Mol. Cell Biol. 15, 482–496 (2014).

55. Tan, J. X. & Finkel, T. Lysosomes in senescence and aging. EMBO Rep. 24, e57265 (2023).

56. Freeman, D. et al. Alpha-synuclein induces lysosomal rupture and Cathepsin dependent reactive oxygen species following endocytosis. PLOS ONE 8, e62143 (2013).

57. Kakuda, K., et al. Lysophagy protects against propagation of α-synuclein aggregation through ruptured lysosomal vesicles. Proc. Natl. Acad. Sci. 121, e2312306120 (2024).

58. Paz, I. et al. Galectin-3, a marker for vacuole lysis by invasive pathogens. Cell. Microbiol. 12, 530–544 (2010).

59. Manecka, D.-L., Vanderperre, B., Fon, E. A. & Durcan, T. M. The neuroprotective role of protein quality control in halting the development of alpha-synuclein pathology. Front. Mol. Neurosci. 10, (2017).

60. Chinta, S. J. et al. Cellular senescence is induced by the environmental neurotoxin paraquat and contributes to neuropathology linked to Parkinson’s disease. Cell Rep. 22, 930–940 (2018).

61. Cohen, J. & Torres, C. Astrocyte senescence: evidence and significance. Aging Cell 18, e12937 (2019).

